# High-Risk Human Papillomavirus E7 Alters Host DNA Methylome and Represses *HLA-E* Expression in Human Keratinocytes

**DOI:** 10.1101/132902

**Authors:** Louis Cicchini, Rachel Z. Blumhagen, Joseph A. Westrich, Mallory E. Meyers, Cody J. Warren, Charlotte Siska, David Raben, Katerina J. Kechris, Dohun Pyeon

## Abstract

Human papillomavirus (HPV) infection distinctly alters methylation patterns in HPV-associated cancer. We have recently reported that HPV E7-dependent promoter hypermethylation leads to downregulation of the chemokine *CXCL14* and suppression of antitumor immune responses. To investigate the extent of gene expression dysregulated by HPV E7-induced DNA methylation, we analyzed parallel global gene expression and DNA methylation using normal immortalized keratinocyte lines, NIKS, NIKS-16, NIKS-18, and NIKS-16ΔE7. We show that expression of the MHC class I genes is downregulated in HPV-positive keratinocytes in an E7-dependent manner. Methylome analysis revealed hypermethylation at a distal CpG island (CGI) near the *HLA-E* gene in NIKS-16 cells compared to either NIKS cells or NIKS-16ΔE7 cells, which lack E7 expression. The *HLA-E* CGI functions as an active promoter element which is dramatically repressed by DNA methylation. HLA-E protein expression on cell surface is downregulated by high-risk HPV16 and HPV18 E7 expression, but not by low-risk HPV6 and HPV11 E7 expression. Conversely, demethylation at the *HLA-E* CGI restores HLA-E protein expression in HPV-positive keratinocytes. Because HLA-E plays an important role in antiviral immunity by regulating natural killer and CD8^+^ T cells, epigenetic downregulation of *HLA-E* by high-risk HPV E7 may contribute to virus-induced immune evasion during HPV persistence.

## INTRODUCTION

Human papillomaviruses (HPV) are small double-stranded DNA viruses with over 180 genotypes that infect mucosal and cutaneous basal epithelia. It has been estimated that up to 80% of sexually active individuals will become infected in their lifetime, making HPV the most common sexually transmitted pathogen^1^. HPVs are classified as high- and low-risk genotypes based on their oncogenic potential^2^. High-risk HPVs are causally associated with ∼5% of human cancers including nearly all cervical cancer (CxCa) and about 25% of head and neck cancer (HNC), making HPV a significant cause of morbidity and mortality worldwide^2,3^. While the majority of primary HPV infections are cleared within two years, ∼10% of infected individuals establish a lifelong persistent infection^4^. Similar studies have revealed that of the genotypes tested, HPV16 is the most likely to persist^5^. Given the propensity of HPV to persist without eliciting a strong immune response, it is very likely that the virus has evolved efficient immune evasion mechanisms.

Dysregulation of hose gene expression is a well-known strategy that viruses frequently employ to evade the host immune response. Of note, Epstein-Barr virus (EBV) hijacks host cell epigenetic machinery to modulate host gene expression^6^. These epigenetic manipulations are considered a hallmark of EBV-induced lymphomas, and persist even after infection is cleared^6,7^. Interestingly, HPV-positive HNC and CxCa progression exhibit distinct changes in host DNA methylation that alter host gene expression^8,9^. In a similar study, HPV-induced cell immortalization corresponded with hypermethylation at several host chromosomal loci including the telomerase subunit *hTERT*^10^. Expression of *hTERT* is increased by promoter hypermethylation which correlates with HPV-associated transformation and cancer progression^11^.

Interestingly, E7 directly binds and activates DNA methyltransferase 1 (DNMT1), leading to a potential epigenetic mechanism of E7-mediated transcriptional modulation^12,13^. Consistently, the HPV E7-DNMT1 complex induces hypermethylation of the tumor suppressor cyclin A1 (*CCNA1*) promoter, an epigenetic marker strongly correlated with HPV-associated malignancy^13,14^. Further, our recent work has revealed that the chemokine *CXCL14* is significantly downregulated by E7-directed promoter hypermethylation^15^. Restoration of *CXCL14* expression in HPV-positive cancer cells prevents tumor formation *in vivo* and increases natural killer (NK) and CD8^+^ T cell populations in the tumor-draining lymph nodes^15^. Downregulation of *CXCL14* is therefore an important immune evasion mechanism employed by HPV E7, allowing for virus persistence. Thus, it is likely that HPV E7 dysregulates other host gene expression by modulating DNA methylation to establish persistent virus infection.

To identify key host factors and pathways altered by HPV-directed DNA methylation in human keratinocytes, we performed parallel global gene expression and DNA methylation analyses. Here, we report that most class I major histocompatibility complex (MHC-I) molecules are transcriptionally downregulated in an E7-dependent manner. Further, non-classical *HLA-E*, which regulates NK and CD8^+^ T cells, is significantly downregulated by E7-mediated hypermethylation in a distal regulatory CpG island (CGI). These results suggest that HPV E7-mediated DNA methylation may modulate host immune responses by downregulating *HLA-E* expression.

## RESULTS

### The HPV oncoprotein E7 drives global gene expression changes in human keratinocytes

To determine gene expression alterations in human keratinocytes by high-risk HPVs, we performed global gene expression analysis in normal immortalized keratinocytes (NIKS) and their derivatives: NIKS-16 and NIKS-18 cells containing episomal HPV16 and HPV18 genomes, respectively. NIKS-16ΔE7 cells containing the HPV16 genome lacking E7 expression^16^ were used to investigate the roles of the HPV oncoprotein E7. Global gene expression profiles of these cells were analyzed using Affymetrix GeneChip Human Genome U133 Plus 2.0 microarrays (GEO accession # GSE83259). Principal component analysis (PCA) of normalized mRNA expression profiles demonstrated that NIKS-16 and NIKS-18 clustered together distinctly from NIKS and NIKS-16ΔE7 cells (**Fig. 1a**). NIKS-16ΔE7 cells, growing slower than the parental NIKS cells, are more morphologically diverse compared to the other NIKS cells tested. These differences may be reflected to high variations between replicates shown by PCA (**Fig. 1a**).

**Figure 1.**
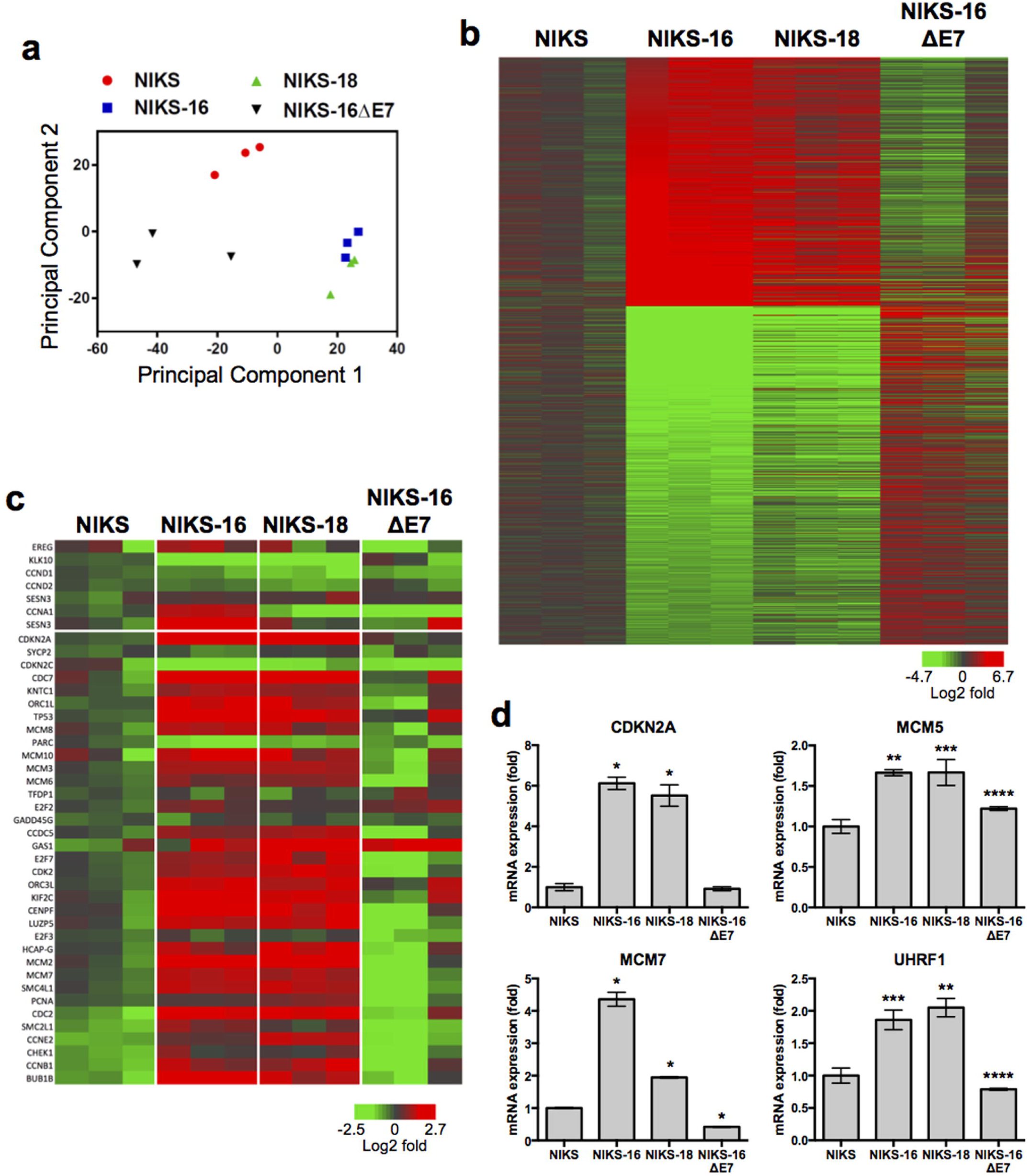
High-risk HPV E7 distinctly alters host gene expression in keratinocytes. Gene expression profiles were assessed by Affymetrix Human Genome U133 Plus 2.0 arrays in triplicate for keratinocytes lines, NIKS, NIKS-16, NIKS-18, and NIKS-16ΔE7, in three different passages. (**a**) Principal component analysis data are shown for each of three replicates of NIKS (red circle), NIKS-16 (blue square), NIKS-18 (green triangle) and NIKS-16ΔE7 (black triangle) cells. (**b**) Log2 fold changes of differentially expressed genes in both NIKS-16 vs. NIKS cells and NIKS-16 vs. NIKS-16ΔE7 cells are shown by heat map (FDR adjusted *p* < 0.05 and a change > 30% magnitude in expression). Probe IDs are listed in **Table S1**. (**c**) Heat map presents log2 fold changes (NIKS-16 vs. NIKS cells and NIKS-16 vs. NIKS-16ΔE7 cells) of dysregulated cell cycle-related genes previously identified in HNC and CxCa patient tissue samples^17^. (**d**) *CDKN2A, MCM5, MCM7*, and *UHRF1* expression levels were determined by RT-qPCR and normalized to β-actin expression levels. Each sample was quadruplicated, and shown are representative of three repeats. Fold changes compared to NIKS cells are plotted. *P*-values were calculated by Student’s *t* test. \**p* < 0.0001, \*\**p* < 0.001, \*\*\**p* < 0.01, \*\*\*\**p* < 0.05.

High-risk HPV infection significantly changes host gene expression patterns including dramatic upregulation of DNA replication- and cell cycle-related gene expression^17,18^. However, the extent of E7-specific gene expression changes has not been fully determined. To define E7-mediated gene expression changes in keratinocytes, we analyzed global gene expression and defined genes up- or downregulated in both comparisons of NIKS-16 vs. NIKS cells and NIKS-16 vs. NIKS-16ΔE7 cells. We identified 625 upregulated and 849 downregulated genes exhibiting a false discovery rate (FDR)-adjusted *p*-value of less than 0.05 for each comparison and a change greater than 30% magnitude in expression (**Fig. 1b, Table S1**). To examine the physiological relevance of HPV-specific gene expression changes, we analyzed the expression patterns of distinct cell cycle-specific genes which were previously identified using CxCa, HNC, and normal patient tissue samples^17^. Consistent with the results from patient tissues, the majority of the cell cycle genes upregulated in HPV-positive cancers were markedly increased in NIKS- 16 and NIKS-18 cells compared to NIKS cells, while none of the cell cycle genes upregulated in HPV-negative HNC tissues were increased in NIKS-16 and NIKS-18 cells (**Fig. 1c**). Interestingly, most of the upregulated genes in NIKS-16 and NIKS-18 cells were not changed or were slightly downregulated in NIKS-16ΔE7 cells compared to NIKS cells. These results indicate that the distinct patterns of cell cycle dysregulation in HPV-positive cancers are largely caused by the HPV oncoprotein E7. Using RT-qPCR, we further validated expression changes of selected genes from **Fig. 1c** (*CDKN2A* and *MCM7*) and previously reported (*UHRF1* and *MCM5*)17,18 (**Fig. 1d**). These results indicate a significant role of the HPV oncoprotein E7 in global gene expression changes during persistent HPV infection in keratinocytes including cell cycle-related genes.

### The HPV oncoprotein E7 downregulates gene expression related to antigen presentation

To understand the biological functions of the identified HPV16 E7-regulated genes (**Table S1**), we performed pathway analysis using Reactome (reactome.org). Consistent with our previous findings^17,18^, the majority of the upregulated genes were involved in cell cycle progression, DNA replication, and DNA repair (**Table S2, Fig. S1a**). In contrast, the pathways of downregulated genes are diverse, suggesting that E7-mediated downregulation of gene expression is more complex than E7-mediated upregulation of gene expression. Interestingly, our analysis revealed that genes involved in antigen presentation, IL1 signaling and extracellular matrix degradation are significantly downregulated in NIKS-16 cells compared to NIKS and NIKS-16ΔE7 cells (**Table S2, Fig. S1b**). Various matrix metalloproteinases and kallikreins were significantly downregulated in NIKS-16 cells compared to NIKS cells and NIKS-16ΔE7 cells (**Table S2, Fig. S2a**).

Importantly, immune response pathways were among the most significantly affected by HPV16 E7 expression (**Figs. S1b, Table S2**). Several genes in antigen presentation (e.g. *HLA-B, HLA-E, SEC31A, ITGAV, CTSL2*, and *RNASEL*) and IL1 signaling (e.g. *IL1B, IL1R1, IL1RN*, and *IL36G*) were significantly altered in NIKS-16 and NIKS-18 cells, but not in NIKS-16ΔE7 cells, compared to NIKS cells (**Fig. S2b and S2c**). Using RT-qPCR, we further validated E7-dependent dysregulation of genes involved in IL1 signaling (**Fig. S2d**). Previous studies have shown that HPV16 E5 disrupts trafficking of MHC-I and -II complexes to the cell surface, and HPV16 E7 downregulates cell surface expression of MHC-I complexes^19–21^. While multiple mechanisms of inhibiting MHC-I surface expression have been observed, HPV-mediated alterations in MHC-I gene expression is poorly understood. Thus, we further assessed expression of all MHC-I α-subunits (*HLA-A, -B, -C, -E, -F,* and *-G*) in the NIKS cell lines. The results showed that except for *HLA-F*, all MHC-I α-subunit gene expression was downregulated in NIKS-16 and NIKS-18 cells compared to NIKS and NIKS-16ΔE7 cells (**Fig. 2a**). To determine any difference in MHC-I gene expression between HPV-positive and HPV-negative cancer tissues, we analyzed the TCGA data of HNC and CxCa obtained from cBioPortal. Interestingly, the expression levels of *HLA-C* and *HLA-E* were significantly lower in HPV-positive HNCs and CxCa than HPV-negative HNCs (**Fig. 2b**). This observation underscores the effect of HPV on HLA gene expression in HPV-associated disease. Unfortunately, due to the presence of numerous splice isoforms for MHC-I genes, we were unable to reliably detect amplicons from MHC-I mRNA transcripts by RT-qPCR. However, all array probesets for *HLA-A*, -*B,* -*C*, -*E* and -*G* detection consistently exhibit significant downregulation in HPV-positive keratinocytes in an E7-dependent fashion (**Fig. 2a**). Overall downregulation of MHC-I genes suggests that HPV16 E7 plays an important role in immune dysregulation of HPV-infected keratinocytes by altering immune cell recognition during early stages of persistent infection.

**Figure 2.**
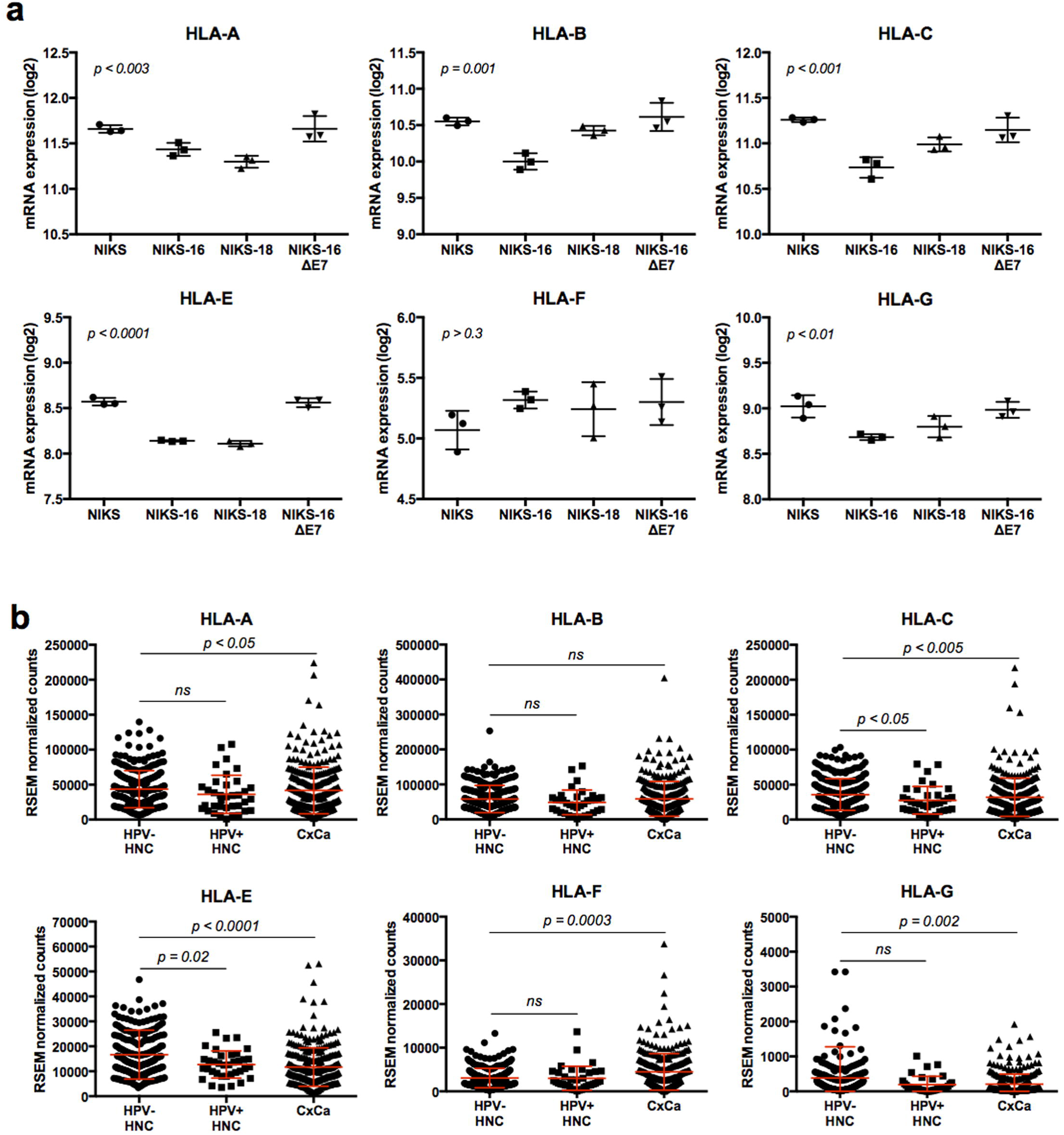
MHC-I gene expression is downregulated in HPV-positive keratinocytes and cancers in an E7-dependent manner. (**a**) Normalized gene expression of *HLA-A, -B, -C, -E, -F*, and *-G* in triplicated NIKS (circle), NIKS-16 (square), NIKS-18 (triangle), and NIKS-16ΔE7 (inverse triangle) cells is shown. Fluorescence intensity (log2) of each replicate is plotted with *p*-values calculated by one-way ANOVA test. (**b**) The RNA-seq RSEM (RNA-seq by expectation maximization) counts of *HLA-A, -B, -C, -E, -F,* and *-G* were obtained from the TCGA data through cBioPortal (cbioportal.org): HPV-negative HNC, *n* = 243; HPV-positive HNC, *n* = 36; CxCa, *n* = 309 (NCI, TCGA, Provisional). Normalized RSEM counts are shown in scatter plots with mean and standard deviation. *P*-values were determined by Mann-Whitney test. *n.s.*, not significant.

### The HPV oncoprotein E7 dysregulates DNA methylation in human keratinocytes

A previous study has shown that HPV infection distinctly modifies the DNA methylation patterns in HNC patients^22^. Additionally, HPV16 E7 protein directly binds to DNMT1 and activates its enzymatic activity^12^. We recently reported that E7-dependent methylation of the *CXCL14* promoter resulted in *CXCL14* downregulation and inhibition of antitumor immune responses^15^. These findings suggest that high-risk HPV E7 is very likely to dysregulate host gene expression by modulating DNA methylation. To investigate the extent of gene expression dysregulated by HPV E7-induced DNA methylation, we analyzed the methylome of NIKS, NIKS-16, NIKS-18, and NIKS-16ΔE7 cell lines in triplicate using Illumina Infinium HumanMethylation450 BeadChip arrays (GEO accession # GSE83261). PCA of methylome showed that each cell type clustered distinctly (**Fig. 3a**). Given that the PCA from our gene expression analysis showed high similarity between NIKS-16 and NIKS-18 cells (**Fig. 1a**), the methylome data may discriminate the molecular patterns of different cell types more precisely than the gene expression data. Nevertheless, the sample-by-sample variations in the triplicates of each cell line are much lower in the methylome data (**Fig. 3a**) than in gene expression data (**Fig. 1a**). This implies that DNA methylation could be a better biomarker than gene expression for early detection of high-risk HPV infection.

**Figure 3.**
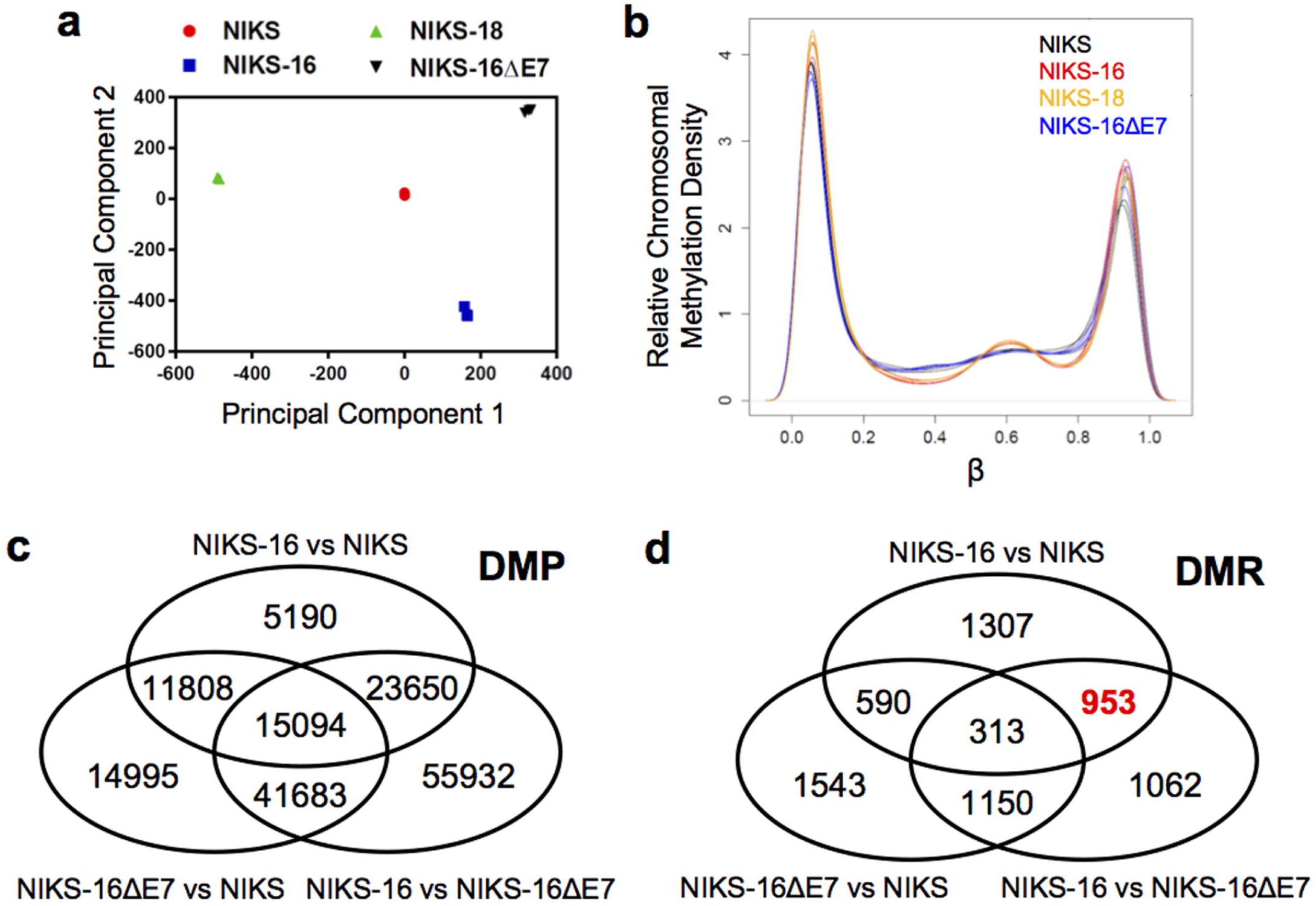
HPV16 E7 alters host genome methylation in keratinocytes. Global DNA methylation profiles in NIKS, NIKS-16, NIKS-18, and NIKS-16ΔE7 cells were analyzed in triplicate using Illumina Infinium HumanMethylation450 BeadChip arrays. (**a**) Principal component analysis data are shown for each replicate of normalized data from NIKS (red circle), NIKS-16 (blue square), NIKS-18 (green triangle) and NIKS-16ΔE7 (black triangle) cells. (**b**) Methylation array data from NIKS (black), NIKS-16 (red), NIKS-18 (orange) and NIKS-16ΔE7 (blue) cells were normalized using SWAN and the relative methylation (β) density across the genome are plotted. β represents the ratio of methylated signal to total signal (methylated + unmethylated) at a given CpG site. β near 0 or 1 indicates no methylation or complete methylation, respectively. Three pairwise comparisons are summarized by Venn diagrams showing the number of overlapping (**c**) differentially methylated positions (DMP, FDR adjusted *p* < 0.05) and (**d**) differentially methylated regions (DMR, permutation *p* < 0.05). DMPs are defined as a single differentially methylated CpG site between groups while DMRs consist of at least two CpG sites separated by no more than 500 bp meeting the cutoff determined by bumphunter algorithm and permutation *p*-value < 0.05.

By assessing the relative methylation density at any given CpG site across the genome, we found that NIKS cells tended to maintain β-values (the ratio of methylated probe intensity vs. the overall intensity) near 0 (0% methylation) or 1 (100% methylation) with uniform distribution between those two peaks on both flanks (**Fig. 3b**, black line). In contrast, both NIKS-16 and NIKS-18 cells exhibited an influx in hemi-methylation near β = 0.6 (**Fig. 3b**, red and orange lines). Interestingly, the methylation pattern of NIKS-16ΔE7 cells was strikingly similar to the methylation pattern of NIKS cells but distinct from the methylation pattern of NIKS-16 and 18 cells (**Fig. 3b**, blue line). These results suggest that E7 alters the global methylation patterns of host genome.

To validate our methylome array data, we assessed DNA methylation status at the *CCNA1* and *TERT* promoter regions that are known to be hypermethylated in HPV-positive cells. Consistent with previous findings, the *CCNA1* and *TERT* promoter regions showed significantly increased methylation (24-28% increase in β, *p* < 0.004) in NIKS-16 cells compared to NIKS cells (**Table S3**)^10,13,14,23^. However, NIKS-18 cells did not show consistent changes in *CCNA1* and *TERT* promoter methylation. Given that the global methylome data showed the distinct DNA methylation patterns between NIKS-16 and NIKS-18 cells, these results suggest that HPV16 and HPV18 may differentially modulate host DNA methylation.

A genome-wide comparison of methylated CpG sites between NIKS and NIKS-16 cells revealed 5,190 differentially methylated positions (DMPs, defined as a single differentially methylated CpG site) and 1,307 differentially methylated regions (DMRs, defined as a cluster of two or more CpG sites, permutation *p*-value < 0.05) (**Fig. 3c and 3d**). To investigate DNA methylation specifically regulated by HPV16 E7, we identified 953 DMRs (red bold in **Fig. 3d**) in a comparison between NIKS-16 to NIKS-16ΔE7 cells and excluded the DMRs found in the NIKS-16ΔE7 to NIKS comparison from the 953 DMRs. Using a more stringent DMR area *p*-value less than 0.01, we identified 56 hypermethylated DMRs (**Table S4**) and 47 hypomethylated DMRs (**Table S4**) that are dependent on HPV16 E7 expression. Interestingly, regional methylation analysis near *HLA-E* identified two significantly hypermethylated DMRs across a total of 5 probed CpG sites (*p* ≤ 0.004, **Table S4**), suggesting that HPV16 E7 may mediate DNA methylation of the *HLA-E* gene.

To determine gene expression regulated by E7-mediated DNA methylation, we identified genes that show gene expression changes (> 30%) with an FDR adjusted *p*-value less than 0.01 and associated DMPs with an FDR adjusted *p*-value less than or equal to 0.005 between NIKS and NIKS-16 cells (**Table S5**). DMPs rather than DMRs were used in this analysis to reduce the possibility of type II errors: the locations of methylation array probes are predicted to be sentinel CpG sites and may not be clustered near additional probes, thus potential true positives may be eliminated from DMR classification. A total of 83 genes showed significant changes of both gene expression and DNA methylation comparing NIKS-16 cells to NIKS cells. This result is consistent with a previous global DNA methylation study assessing epigenetic changes directed by EBV, showing that most DMPs are silent and relatively a small number of them contributed to gene expression changes^7^. Our results suggest that E7-mediated DNA methylation regulates gene expression of a subset of host genes. Interestingly, *HLA-E* shows a significant decrease in gene expression and increase in DNA methylation (**Figs. 2 and 4**). Consistent with the *HLA-E* gene expression results, the comparison between NIKS-16 and NIKS-16ΔE7 cells showed a significant decrease in DNA methylation at *HLA-E* (24%) in NIKS-16ΔE7 cells compared to NIKS-16 cells (**Table S5**). These results suggest that downregulation of *HLA-E* gene expression shown in **Fig. 2a** is likely caused by HPV16 E7-mediated DNA methylation.

### DNA methylation of the *HLA-E* CGI is significantly increased by the HPV oncoprotein E7

Our methylome data consistently showed that DNA methylation of *HLA-E* was significantly increased in NIKS-16 cells compared to NIKS cells in an E7-dependent manner (**Table S5**). Unexpectedly, the *HLA-E* CGI containing the identified DMR is ∼23,000 bases upstream of the *HLA-E* open reading frame (ORF), potentiating its functional role as an enhancer or distal promoter element. Our results from scanning the *HLA-E* CGI for DNA methylation showed a dramatic increase (∼50%) near the 3’ region of the CGI (**Fig. 4a**). To validate DNA methylation in the *HLA-E* CGI, we performed methylation-specific PCR (MSP) using primer sets specific to the *HLA-E* DMR. Consistent with the methylome data, the 3’ region of the *HLA-E* CGI was highly methylated in NIKS-16 cells compared to NIKS cells, but the *HLA-E* CGI methylation dramatically decreased in NIKS-16ΔE7 cells (**Fig. 4b**). The DNA methylation of the *HLA-E* CGI is highly correlated with the decrease of *HLA-E* expression in NIKS-16 cells, but not in NIKS-16ΔE7 cells (**Fig. 2a**). These results suggest that *HLA-E* gene expression may be downregulated by HPV16 E7-mediated DNA methylation.

**Figure 4.**
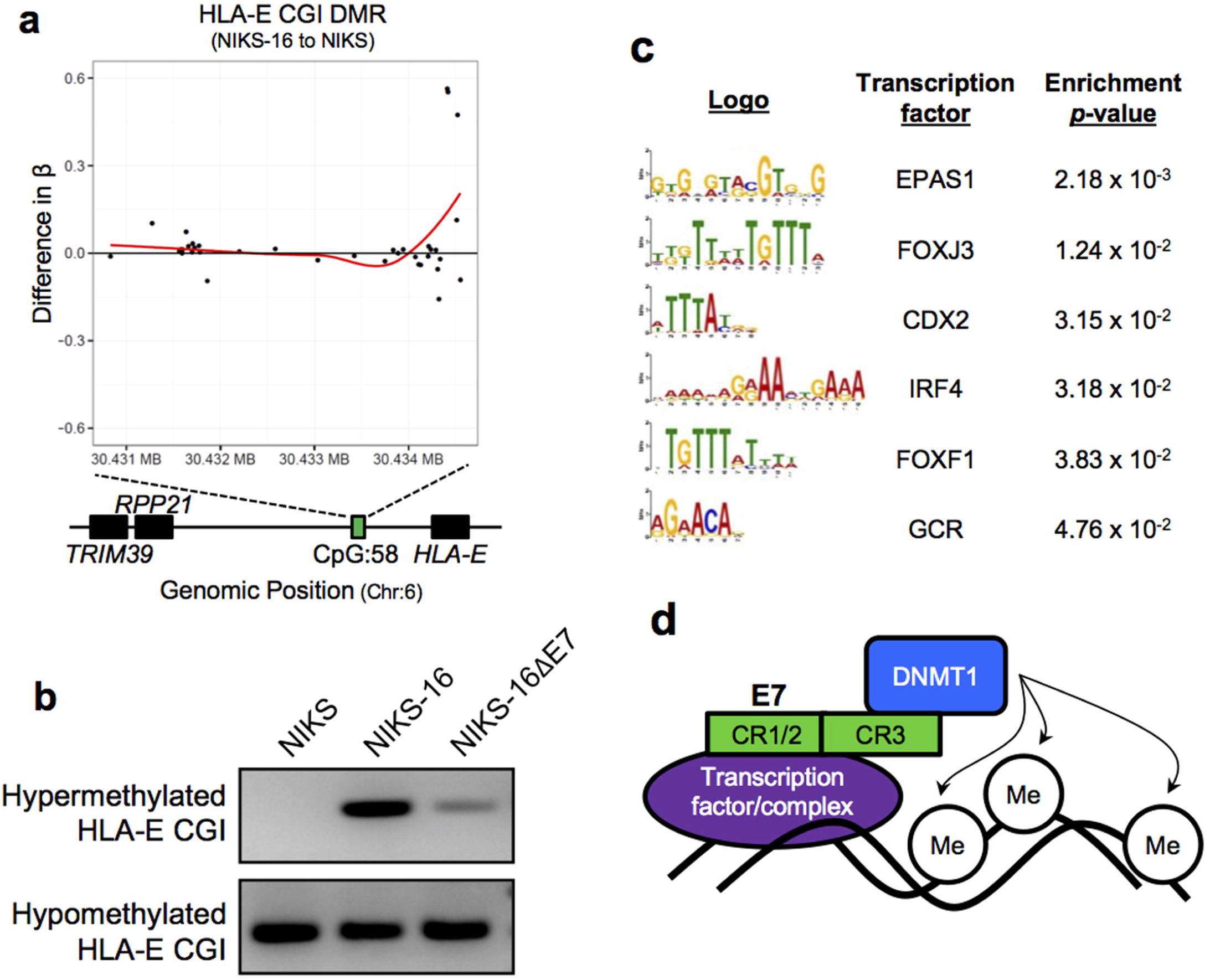
HPV16 E7 is necessary for hypermethylation at a distal *HLA-E* CpG island. (**a**) The difference in methylation (β) of all probed CpG dinucleotides in the *HLA-E* CpG island (CGI, chr6:30,434,030-30,434,730) between NIKS and NIKS-16 cells is shown. Positive and negative β indicates increased or decreased methylation in NIKS-16 cells compared to NIKS cells, respectively. The red line represents a locally weighted scatter plot (LOESS) regression curve, showing the trend of differences in Beta values (Y) along the genomic position (X). (**b**) Methylation specific PCR (MSP) products were separated in 2% agarose gel to evaluate the methylation status of the *HLA-E* CGI using bisulfite-converted gDNA from NIKS, NIKS-16, and NIKS-16ΔE7 cells and primers listed in **Table S7**. (**c**) Sequence logos of enriched transcription factor (TF) binding motifs are shown. 100 bp regions flanking E7 sensitive DMRs (comparing NIKS to NIKS-16 cells, *p* < 0.04, count 185) were assessed for enrichment of TF binding motifs using MEME Suite software and nucleotide frequencies in the submitted sequences as background (enrichment *p* < 0.05). (**d**) Schematic diagram of the potential mechanism of targeted E7-induced DNA methylation. E7 binds transcription factors (or complexes) through its CR1/2 domain and DNMT1 through its CR3 domain, leading to hypermethylation near specific transcription factor binding motifs.

HPV16 E7 directly binds and activates DNMT1 through its CR3 zinc finger binding domain, providing a potential mechanism of E7-induced DNA methylation^12^. However, it is unclear how E7-mediated DNA methylation targets specific regions in the genome. We hypothesized that the regions near specific transcription factor (TF) binding sites are targeted by the E7-DNMT1 complex to direct DNA methylation. Supporting our hypothesis, a previous study revealed that HPV16 E7 recruits histone deacetylases (HDACs) to IRF-1 regulatory promoter complexes thereby directing histone deacetylation to silence IRF-1 responsive genes^24^. To test our hypothesis, we compiled a list of hypermethylated DMRs, filtered as described above for E7-dependent hypermethylation (*p* < 0.04, count 185) and submitted DMRs with 100 bp of flanking sequence to MEME suite for enrichment analysis of TF binding motifs. Interestingly, E7-dependent hypermethylated DMRs showed enrichment of EPAS1, FOXJ3, CDX2, IRF4, FOXF1, and glucocorticoid receptor (GCR) TF binding sites (**Fig. 4c**). Enrichment of IRF4, FOXF1 and GCR motifs imply that E7-mediated DNA methylation may be directed to TF binding motifs near immunoregulatory and developmental genes^25–28^. Consistently, scanning the *HLA-E* CGI (containing the identified DMR) for TF binging sites identified a GCR consensus binding motif, AGAACA (**Fig. 4c**). Previous studies have shown that GCR is involved in suppression of MHC-I27 and MHC-II expression^29^. Enrichment of methylation near specific TF binding sites implies that HPV E7 may direct DNA methylation by recruiting DNMT1 methyltransferase to specific promoter elements through interactions with TFs (**Fig. 4d**). Further analysis revealed that the *HLA-E* CGI contains sites for DNase I hypersensitivity and acetylated H3K27 histone markers, both indicative of active regulatory elements. Additionally, small noncoding RNAs (ncRNAs) are transcribed from 3’ of the *HLA-E* CGI (**Fig. S3**). MicroRNA target prediction analysis of these ncRNAs revealed 65 putative target cellular mRNAs (**Table S6**), including histocompatibility 13 (*HM13*), which is involved in loading peptides onto HLA-E. HLA-E surface expression and stabilization require antigen binding, suggesting a potential mechanism of downregulating HLA-E surface expression^30^. Taken together, the *HLA-E* CGI appears to be an active site for transcription of ncRNAs and other regulatory elements which may have direct or indirect effects on *HLA-E* expression.

### The promoter activity of the *HLA-E* CGI is repressed by DNA methylation

To assess the transcriptional regulation by DNA methylation at the *HLA-E* CGI, we employed a promoter reporter assay using a CpG-free firefly luciferase expression vector, pCpGL-Basic^31^. We first determined the promoter and/or enhancer activity of the *HLA-E* CGI by cloning the *HLA-E* CGI (hg19, chr6:30,434,030-30,434,730) into the pCpGL-Basic vector. pCpGL-HLAE-CGI constructs were prepared to test the promoter activity of the *HLA-E* CGI in forward and reverse orientations (pCpGL-HLAE-Fwd and pCpGL-HLAE-Rev). Additionally, we tested the enhancer activity of the *HLA-E* CGI in combination with a downstream EF1α promoter (pCpGL-HLAE-Fwd-EF1α and pCpGL-HLAE-Rev-EF1α) (**Fig. 5a**). Each construct was transfected into 293FT cells along with a *Renilla* luciferase vector as a transfection control. Promoter activity was determined by relative luciferase activity. Our results revealed the strong promoter activity of the *HLA-E* CGI, showing near 200-fold and 130-fold increases in luciferase activity by insertion of the *HLA-E* CGI in forward or reverse orientations, respectively (**Fig. 5b**).

**Figure 5.**
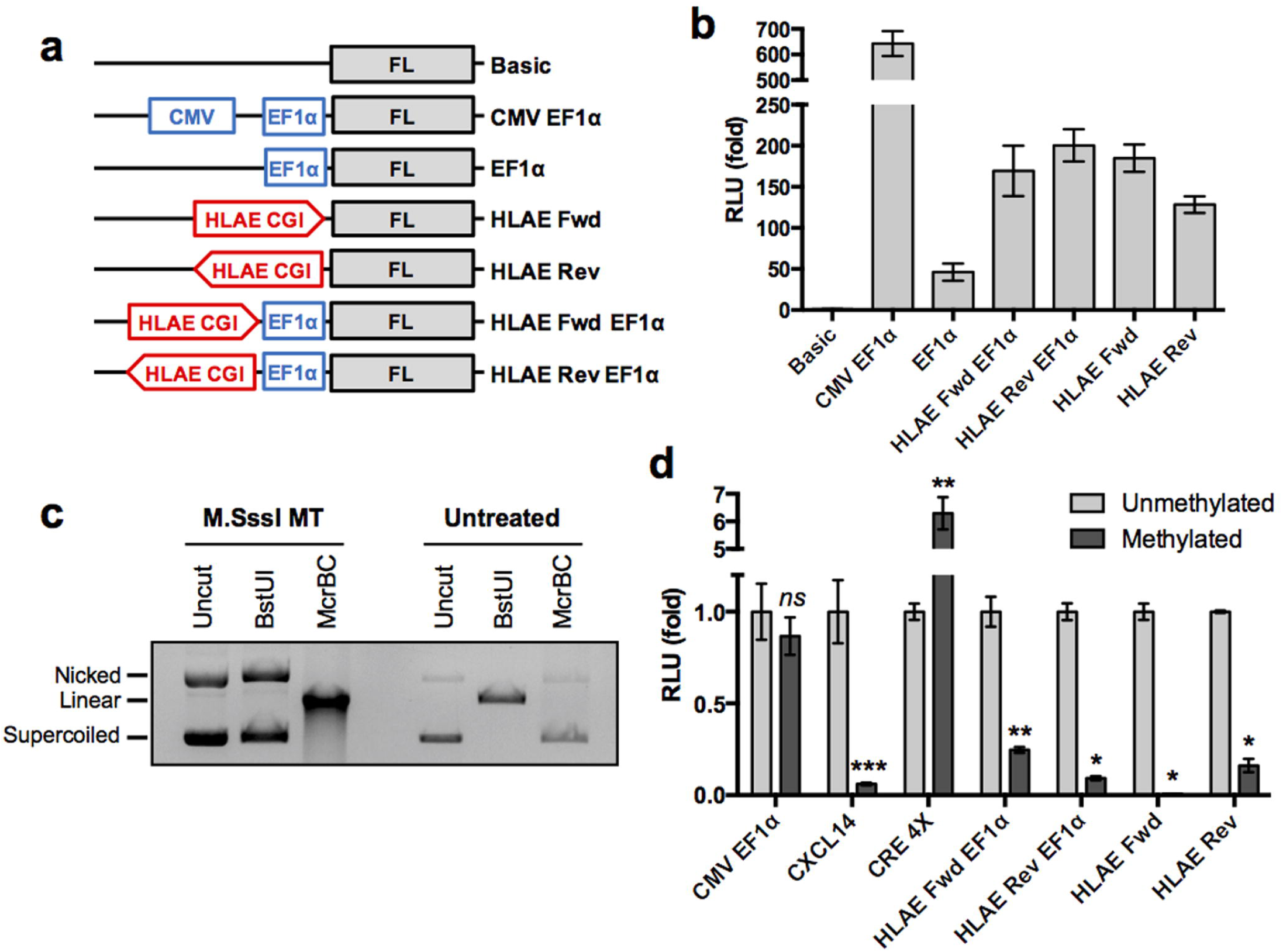
Promoter activity of the *HLA-E* CGI is repressed by DNA methylation. (**a**) Schematic representation of pCpGL plasmid constructs. The *HLA-E* CGI (HLAE CGI) was directionally cloned into pCpGL constructs indicated by an arrow head at the 3’ end. (**b**) 293FT cells were transfected with indicated pCpGL constructs (panel **a**) along with a *Renilla* luciferase (RL) plasmid as a transfection control. Luciferase activity was measured 24 hours post transfection using the Dual Luciferase Reporter Assay (Promega). Representative data of three independent experiments is shown as a fold ratio (FL/RL) of relative light units (RLU) from quadruplicates. (**c**) The pCpGL plasmid containing the *CXCL14* promoter element was incubated with M.SssI methyltransferase (MT) or buffer only (untreated). Samples were subsequently treated with buffer only (uncut), BstUI or McrBC methylation-sensitive endonucleases. Samples were separated in 0.7% agarose gel to verify *in vitro* methylation. (**d**) 293FT cells were transfected with M.SssI MT-treated (methylated) or buffer only control (unmethylated) pCpGL reporter constructs (described in panel **a**) along with an RL plasmid as a transfection control. Luciferase activity was assessed. Fold changes to the unmethylated reporter constructs are plotted. Shown are representative data of four to five repeats. *P*-values were calculated by Student’s *t* test. \**p* < 0.0005, \*\**p* < 0.005, \*\*\**p* = 0.01.

We next tested if hypermethylation in the *HLA-E* CGI represses its promoter activity using *in vitro* DNA methylation. The pCpGL reporter constructs were methylated *in vitro* using the M.SssI CpG methyltransferase. To verify successful DNA methylation of the pCpGL reporter constructs, methylated and unmethylated plasmids were digested with restriction enzymes BstUI and McrBC, which cut only unmethylated and methylated CpG motifs, respectively (**Fig. 5c**). Each methylated and unmethylated plasmid was transfected into 293FT cells and relative luciferase activity was measured. The CpG-free pCpGL-CMV-EF1α plasmid (unaffected by CpG methylation) was used as a negative control^31^ and the pCpGL-CXCL^14^ promoter (repressed by CpG methylation) plasmid and pCpGL-CRE4X (activated by CpG methylation) plasmid were used as positive controls^15,32^. Interestingly, *in vitro* DNA methylation dramatically decreased the luciferase activity of all *HLA-E* CGI containing reporter plasmids (**Fig. 5d**). As expected, while the luciferase activity of pCpGL-CMV-EF1α was unchanged, the luciferase activity of pCpGL-CXCL14 and pCpGL-CRE4X and were significantly decreased and increased by *in vitro* DNA methylation, respectively (**Fig. 5d**). These results suggest that *HLA-E* expression is negatively regulated by DNA methylation in the *HLA-E* CGI.

### HLA-E protein expression is downregulated by high-risk HPV E7, but not by low-risk HPV E7

To determine if HLA-E protein levels are also decreased by E7, cell surface expression of HLA-E proteins was determined in NIKS, NIKS-16, and NIKS-18 cells by flow cytometry. Gating for flow cytometry and staining controls are shown in **Figs. S4 and S5**, respectively. As previous studies have shown that surface expression of MHC-I molecules is frequently downregulated in HPV-positive cells and tissues^33^, we also examined HLA-B/C expression in NIKS, NIKS-16, and NIKS-18 cells. Consistent with our mRNA expression data, protein expression of HLA-E as well as HLA-B/C was dramatically decreased in NIKS-16 and NIKS-18 cells compared to NIKS cells (**Fig. 6a and 6b**). As shown above, HPV16 E7 expression is necessary for *HLA-E* downregulation (**Fig. 2a**). To test if expression of high-risk E7 is sufficient for HLA-E downregulation, we generated stable NIKS cell lines expressing E7 from high-risk HPV genotypes (16 and 18) and low-risk HPV genotypes (6 and 11). E7 expression in each NIKS cell line was validated by RT-PCR (**Fig. S6**), as antibodies detecting E7 from different genotypes are not available. Interestingly, high-risk HPV16 E7 or HPV18 E7 expression was sufficient for decrease of HLA-E proteins on NIKS cells, while low-risk HPV6 E7 or HPV11 E7 expression rather increased HLA-E expression on NIKS cells (**Fig. 6c and 6d**). In contrast, low-risk E7 expression moderately decreased HLA-B/C expression compared to substantial downregulation of HLA-B/C by high-risk E7 expression (**Fig. 6e and 6f**). These results highlight the distinct functions of high-risk and low-risk E7 proteins in dysregulation of MHC-I expression.

**Figure 6.**
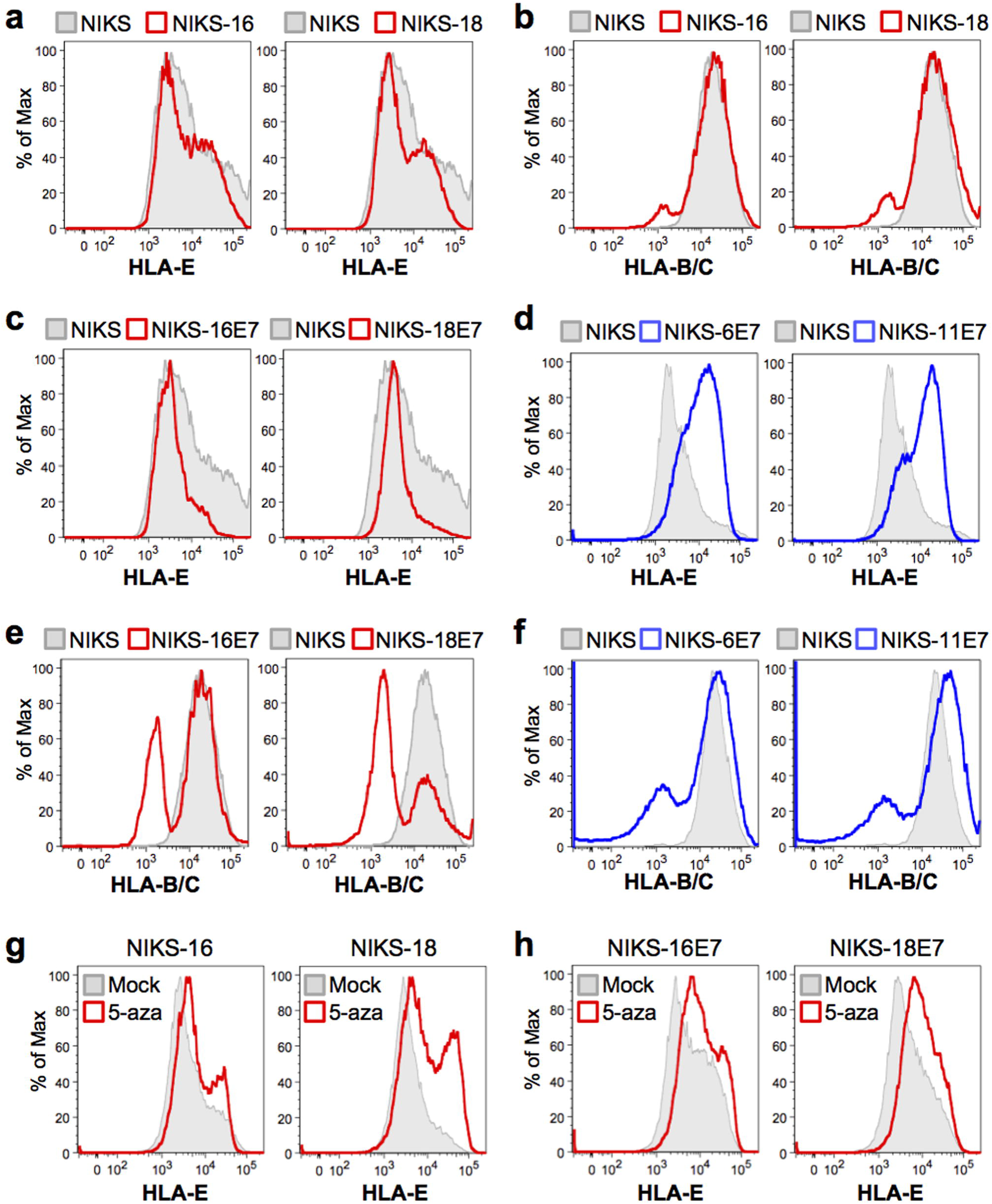
High-risk HPV E7, but not low-risk HPV E7, is sufficient for downregulation of *HLA-E* protein expression in keratinocytes. NIKS and NIKS derivative cells were dissociated to single cell populations using citric saline buffer, fixed in 4% paraformaldehyde, incubated with HLA-E or HLA-B/C antibodies, and assessed by flow cytometry as described in Supplementary Methods. HLA-E (**a**) and HLA-B/C (**b**) protein expression in NIKS (grey) and NIKS-16 (red) cells. (**c-f, and h**) NIKS cells stably expressing E7 from HPV6, 11, 16, and 18 (NIKS-6E7, NIKS-11E7, NIKS-16E7, and NIKS-18E7, respectively) were generated by lentiviral transduction followed by puromycin selection. HLA-E (**c and d**) and HLA-B/C (**e and f**) expression in NIKS cells (grey) and NIKS cells expressing high-risk (HPV16 and 18, red) or low-risk (HPV6 and 11, blue) E7 was analyzed by flow cytometry. HLA-E expression in NIKS-16 cells mock treated or treated with 5 μM 5-aza for five days was assessed by flow cytometry (**g and h**). Shown are representative data of three repeats.

As we showed that demethylation at the *HLA-E* CGI restored *HLA-E* gene expression, we treated NIKS-16 cells with the demethylating agent, 5-aza-2’-deoxycytidine (5-aza). Interestingly, 5-aza treatment dramatically restored HLA-E protein expression on NIKS-16 and NIKS-18 cells (**Fig. 6g and 6h**). Demethylation at the *HLA-E* CGI with 5-aza treatment in NIKS-16 cells was verified by MSP (**Fig. S7**). Taken together, our findings suggest that HLA-E expression is downregulated by the HPV oncoprotein E7-mediated DNA methylation and can be restored by treatment with a demethylating agent.

## DISCUSSION

Our previous global gene expression studies using CxCa and HNC patient tissue samples have shown that expression of cell cycle-related genes is highly upregulated in HPV-positive cancers compared to normal tissue and HPV-negative HNC^17,18^. Additionally, other studies have shown that HPV alters global DNA methylation as a mechanism to silence host gene expression using genetically dissimilar HNC tissues and cell lines^22,34^. In parallel gene expression and methylome analyses using homogeneous keratinocytes, we found that the *HLA-E* CGI is hypermethylated in HPV-positive cells in an E7-dependent manner, correlating with downregulation of *HLA-E* expression. Previous studies have shown that DNA hypermethylation is an effective mechanism of repressing MHC-I and -II gene expression, which is reversed by methylation inhibitors35,36; however, any effect of HPV on *HLA-E* expression was unknown.

MHC expression is modulated by various cellular mechanisms. For example, the *HLA-G* promoter contains a series of *cis* regulatory elements that govern tissue-specific expression^37^. Distal promoter elements are scattered throughout the MHC gene locus and activate transcription of ncRNAs that may regulate expression of the MHC genes^38^. One distal regulatory element has been characterized 25 kb upstream of the *HLA-DRA* promoter^39^. Similarly, we here report that a distal CGI located 23 kb upstream of the *HLA-E* ORF exhibits strong promoter activity. We showed that the *HLA-E* CGI remains hypomethylated in normal (NIKS) cells (β = 12.4%), but hypermethylated in NIKS-16 cells (change in β >50%) and its methylation is linked to *HLA-E* downregulation. Thus, it is possible that HPV E7-induced DNA methylation silences promoter activity of the distal *HLA-E* CGI, or silences expression of a regulatory ncRNA which may modulate *HLA-E* regulatory elements, as previously hypothesized^38^. Additionally, DNase I hypersensitivity (indicative of a relaxed chromatin structure), H3K27 acetylation (a histone marker synonymous with active transcription)^40^, conserved TF binding sites, and ncRNAs from this region indicate that the *HLA-E* CGI is an active regulatory region for *HLA-E* expression (**Fig. 4c, Fig. S3**).

It has been suggested that HPV E7 inhibition of STAT1 activation represses *TAP1* transcription, which leads to a decrease of surface expression of MHC-I molecules^41^. However, the *TAP1* mRNA levels were not changed in either NIKS-16 or NIKS-18 cells compared to NIKS and NIKS16ΔE7 cells (data not shown). This observation suggests that downregulation of HLA-B/C protein expression in NIKS-16 and NIKS-18 cells might not be mediated by E7-induced TAP1 downregulation. Here, our study showed that *HLA-E* expression is regulated by DNA methylation and is restored by treatment of a demethylating agent. In contrast, the *HLA-B/C* gene regions showed no significant changes in DNA methylation induced by HPV16 or HPV18. Thus, *HLA-B* and -*C* downregulation is likely caused by HPV E7, or is mediated by other unknown mechanisms. Interestingly, the MHC-I transactivator NLRC5, which is necessary for MHC-I gene expression and repressed by DNA methylation in various cancers42, was downregulated in NIKS-16 and NIKS-18 cells when compared to NIKS and NIKS-16ΔE7 cells (**Table S1**). This may in part explain E7-dependent downregulation of HLA-B/C expression, while further studies are essential to fully understand the mechanism.

*HLA-E* is constitutively expressed in various tissues and at different stages in development, suggesting tight spatial-temporal regulation of expression^43^. We observed that HLA-E expression in NIKS cells is not homogeneous (**Fig. 6**). This may be due to heterogeneous populations and/or the nature of the co-culture model with feeders wherein NIKS cells propagate in islands with visually distinct morphologies depending on their relative position to other NIKS cells. It would be interesting, therefore, to further explore HLA-E expression in distinct layers of three-dimensional tissue, which may also dictate HLA-E functionality.

HLA-E regulates NK and T cells through direct contact with surface receptors. The NK cell inhibitory receptor NKG2A and the activating receptor NKG2C were initially found to bind HLA-E on the cell surface^44^. HLA-E presentation of MHC leader peptides to NK cells leads to NKG2A-mediated inhibition of NK effector function^44,45^. Interestingly, HLA-E presentation of stress-inducible heat shock protein 60 peptides interferes with NKG2A recognition, leading to NK cell-mediated killing of stressed cells^46^. In addition to regulating NK cells, HLA-E also acts on CD8^+^ T and natural killer T (NKT) cells to modulate adaptive immune responses^47,48^. HLA-E activates or inhibits NK and CD8^+^ T cell functions by presenting a narrow range of viral and bacterial antigens^49^. Of note, a viral-derived peptide presented by HLA-E on HIV-1-infected CD4+ T cells activates NK cells and induces cytolysis of virus-infected cells^50^. In contrast, a viral peptide presented by HLA-E on the surface of hepatitis C virus-infected cells inactivates NK cells but elicits HLA-E-restricted CD8^+^ T cell responses^51^. These findings imply that HLA-E has the potential to present HPV-derived peptides to NK or CD8^+^ T cells, which may lead to lysis of the HPV-infected cells. Interestingly, we have previously found that high-risk HPV E7 significantly reduces NK and CD8^+^ T cell infiltration into the HPV-positive tumor microenvironment through *CXCL14* downregulation by its promoter hypermethylation^15^. Together, these results suggest that HPV evade antiviral NK and CD8^+^ T cell activity by dysregulating DNA methylation.

In contrast, *HLA-E* is upregulated in several cancers, including HNC, CxCa, breast, rectal, colon, and ovarian cancers^8^. Accordingly, it has been speculated that the high levels of HLA-E may inhibit NK cell activation caused by downregulation of classical MHC-I expression on cancer cells. The better survival rate of ovarian cancer patients with infiltrating CD8^+^ T cells disappears when NKG2A signaling is activated by high *HLA-E* expression^52^. *HLA-E* expression is also linked to poor clinical outcome and low overall survival in breast and colon cancer patients^53,54^. Further, knockdown of *HLA-E* expression enables NKG2D-mediated lysis of glioma cells by NK cells^55^. Our results showing that high-risk HPV E7 downregulates *HLA-E* expression imply dual roles of *HLA-E* that induces antiviral immunity in normal cells but suppresses antitumor immunity in cancer cells.

Indeed, previous studies have shown that HLA-E plays an important role in antiviral immune responses. HLA-E presentation of viral peptides elicits cytotoxic responses of NK and CD8^+^ T cells that kill virus-infected cells^50,51,56^. We report here that high-risk HPV E7 significantly downregulates *HLA-E* expression in keratinocytes, while low-risk HPV E7 increases *HLA-*E expression. The epigenetic repression of *HLA-E* expression by E7 suggests a previously undescribed immune evasion mechanism employed by high-risk E7, but not by low-risk E7. Additionally, HLA-E expression can be restored through treatment with the demethylating agent, 5-aza. This may provide a new therapeutic approach to treat HPV-positive lesions by activating the HLA-E mediated antitumor immune responses of NK and CD8^+^ T cells.

## METHODS

### Cell Culture

Human keratinocytes NIKS^57^, NIKS-16, NIKS-18^58^, and NIKS-16ΔE7^16^ cells were co-cultured with NIH 3T3 feeder cells in E-complete medium as previously described^17^. NIKS-6E7, NIKS-11E7, NIKS-16E7, and NIKS-18E7 cell lines were generated by lentiviral transduction and puromycin selection. HeLa cells were obtained from ATCC and cultured in DMEM supplemented with 10% FBS according to the manufacturer’s recommendations.

### Genome-wide Expression and DNA Methylation Arrays

For gene expression analysis, total RNA was extracted from NIKS, NIKS-16, NIKS-18, and NIKS-16ΔE7 cells using the RNeasy kit (Qiagen) and hybridized to Affymetrix Human Genome U133 Plus 2.0 Array chips as previously described^17^. For methylome analysis, genomic DNA (gDNA) was isolated from NIKS, NIKS-16, NIKS-18, and NIKS-16ΔE7 cells using the DNeasy kit (Qiagen). Bisulfite-converted gDNA was prepared using the EZ DNA Methylation Kit (Zymo Research) and assessed using Illumina Infinium HumanMethylation450 BeadChip Kits according to the manufacturer’s protocol. Data processing methods are discussed in supplemental material.

### Methylation Specific PCR (MSP) and *in vitro* DNA Methylation

Methylation of the *HLA-E* CGI was analyzed by MSP. gDNA was extracted from NIKS, NIKS-16, and NIKS-16ΔE7 cells using DNeasy Blood & Tissue Kit (Qiagen). 500 ng of the gDNA was used in each bisulfite reaction using the EZ DNA Methylation Kit (Zymo Research) according to the manufacturer’s instruction. MSP was performed with primers described in **Table S7** and validated using methylated or unmethylated control gDNA. Control DNA was generated by *in vitro* DNA methylation of gDNA extracted from W12E, W12G, and W12GPXY cells using McrBC endonuclease *or* M.SssI CpG methyltransferase followed by BstUI digestion (New England Biolabs). *In vitro* DNA methylation was performed using the M.SssI CpG methyltransferase and methylation efficiency was validated by McrBC or BStUI digestion. For DNA demethylation, NIKS-16 cells were treated daily with 5 μM 5-aza-2’-deoxycytidine (5-aza) for five days.

### Plasmids and Lentiviral Constructs

The pCpGL-Basic luciferase reporter vector was a gift from Michael Rehli (University of Regensburg, Germany)^31^. The four repeats of the cAMP-response element (CRE4X) were synthesized as oligonucleotides and cloned into pCpGL-Basic using BamHI and NcoI^32^. The *CXCL14* promoter was cloned into pCpGL-Basic using specific primers (**Table S7**) between the BamHI and NcoI sites. A new multiple cloning site (MCS) was introduced into the pCpGL-Basic vector in place of the CMV enhancer by PCR-mediated mutagenesis, and the *HLA-E* CGI was directionally cloned in using specific primers (**Table S7**). For generation of E7-expressing NIKS cells, the HPV E7 genes were obtained from Joe Mymryk (University of Western Ontario, Canada) and cloned into lentiviral expression vectors (pCDH-CMV-MCS-EF1-Puro, System Biosciences) between the XbaI and BamHI sites using PCR with specific primers (**Table S7**).

### Microarray data accession number

The microarray data of gene expression and DNA methylation are accessible in the NCBI GEO database under accession numbers GSE83259 and GSE83261, respectively.

## ACKNOWLEDGMENTS

We thank Michael Rehli for providing the pCpGL vectors, Joe Mymryk for providing HPV6 and 11 E7 constructs, and the Genomics and Microarray Core for providing gene expression and methylome analyses. We also thank Ivana Yang, James Hagman, Lauren Vanderlinden, and members of the Pyeon laboratory for useful comments and suggestions.

## AUTHOR CONTRIBUTIONS

L.C., R.Z.B., M.E.M., J.A.W., K.J.K., and D.P. conceptualized and designed experiments. L.C., R.Z.B., M.E.M., J.A.W., C.J.W., C.S., and D.P. analyzed and interpreted data. L.C. and D.P. drafted the manuscript. L.C., R.Z.B., J.A.W., D.R., K.J.K, and D.P revised and critically reviewed the manuscript.

